# *Enterococcus faecalis* alters antibiotic susceptibility in *Pseudomonas aeruginosa* mixed species biofilms

**DOI:** 10.1101/2025.11.24.690246

**Authors:** Caleb M Anderson, Yves Mattenberger, Ana Parga, Heidi Portalier, Casandra Ai Zhu Tan, Maria Esteban Henao, Patrick H Viollier, Kimberly A Kline

**Author notes:** correspondence to Kimberly A Kline, and Patrick H Viollier.

## Abstract

Bacterial infections often occur in polymicrobial biofilms where nutrient limitation and interspecies interactions can profoundly shape microbial physiology. *Enterococcus faecalis* can antagonize *Pseudomonas aeruginosa* growth under conditions of iron limitation, a known host defense mechanism. We report here that this growth antagonism uncovers surviving *P. aeruginosa* cells capable of surviving antibiotic challenge, including ampicillin, cefepime, and ciprofloxacin, when grown in iron-restricted biofilms with *E. faecalis*. Transcriptomic profiling of *P. aeruginosa* revealed a distinctive response characterized by broad downregulation of biosynthetic, metabolic, and virulence pathways, alongside selective induction of membrane remodeling proteins, transport systems, and biofilm-associated genes. Induction of *arnT* in *P. aeruginosa*, required for lipid A modification, correlated with enhanced antibiotic survival to ampicillin, cefepime, and ciprofloxacin. Additionally, the diguanylate cyclase SiaD and efflux transporter MfsC in *P. aeruginosa* were implicated in decreased antibiotic susceptibility to the same antibiotics above. This transcriptional response was unique to the dual stress of iron deprivation and microbial competition with *E. faecalis*, illustrating how interspecies interactions can simultaneously inhibit and protect *P. aeruginosa*, shedding light on potential persistence mechanisms in iron-limited polymicrobial environments.

**IMPORTANCE:** This study addresses antibiotic susceptibility in *Pseudomonas aeruginosa*, a major opportunistic ESKAPE pathogen, within polymicrobial biofilms and under host-relevant iron-restricted conditions. Polymicrobial biofilm-associated infections are notoriously difficult to treat due to complex interspecies interactions and increased antibiotic resistance. We demonstrate that *Enterococcus faecalis* not only antagonizes *P. aeruginosa* growth under iron limitation but also induces a unique transcriptional profile enhancing *P. aeruginosa* survival during antibiotic challenge. This shift involves broad transcriptional reprogramming in *P. aeruginosa*, characterized by global metabolic downregulation and activation of envelope remodeling pathways, including the *arn* operon. These findings reveal how interspecies interactions under iron stress can both suppress and protect bacterial pathogens and underscore the importance of considering community context in treatment strategies for persistent infections.

## INTRODUCTION

Bacteria typically exist within complex communities where interspecies interactions play a critical role in survival, especially under stressful conditions such as nutrient limitation or antimicrobial pressure (1–14). Understanding these interactions is crucial for elucidating mechanisms of microbial persistence, especially in the context of polymicrobial infections (1, 6, 12, 13, 15). Among the organisms frequently implicated in polymicrobial and biofilm-associated infections are *Pseudomonas aeruginosa* and *Enterococcus faecalis*, opportunistic pathogens that exhibit robust resilience to a broad spectrum of antibiotics (2, 16–18). While each organism is independently capable of causing diverse infections, their frequent co-isolation in polymicrobial biofilm-associated infections, including chronic wound infections, periodontitis, and urinary tract infections, supports more careful study of their interspecies interactions in these environments (3, 19–28).

Interactions between *P. aeruginosa* and *E. faecalis* profoundly alter their individual behavior depending on environmental context (29–33). During co-infection in pyelonephritis, the presence of *E. faecalis* increases *P. aeruginosa* tolerance to β-lactam antibiotics, suggesting a cooperative effect under specific conditions (29). Conversely, under iron-restricted conditions, *E. faecalis* antagonizes *P. aeruginosa* growth, reducing its survival to approximately 0.02% in *in vitro* biofilm assays (33). This antagonism is driven by a combination of competition for iron and environmental acidification caused by *E. faecalis* lactic acid production, illustrating a shift from synergism to antagonism under iron stress (33). Iron is an essential nutrient for bacterial growth and is tightly sequestered in the host as a component of nutritional immunity (34–36). To overcome this limitation, pathogens employ mechanisms such as siderophore production and specialized iron uptake systems adapted to their infection niche (8, 10, 36–38). Under iron-limited conditions, interspecies interactions can exert profound and sometimes unexpected effects on bacterial physiology. We previously showed that *E. faecalis* enhances *E. coli* survival under iron deprivation by exporting L-ornithine, which induces enterobactin synthesis in *E. coli* (39). This relationship contrasts with the antagonistic interaction observed *in vitro* between *P. aeruginosa* and *E. faecalis* under iron starvation, an effect driven by both environmental acidification and competition for iron (33). The molecular responses of *P. aeruginosa* to the competition *E. faecalis* under these conditions remain unclear.

In this study, we sought to elucidate the consequences of *E. faecalis*-*P. aeruginosa* interactions, focusing on how these interactions influence antibiotic susceptibility under iron limited conditions. Our results reveal that although *E. faecalis* antagonizes *P. aeruginosa* growth, it significantly enhances broad-spectrum antibiotic survival among the remaining *P. aeruginosa* population. Growth with *E. faecalis* under iron restriction reshapes the transcriptome of *P. aeruginosa*, resulting in broad downregulation of essential metabolic, regulatory, stress response, and virulence pathways, coupled with selective upregulation of cell envelope remodeling genes. Among these, we identified upregulation of *arnT*, which mediates the transfer of L-Ara4N to lipid A, a modification known to reduce susceptibility to cationic antimicrobials and thereby confer increased antibiotic tolerance in *P. aeruginosa* (40–43). We show that *arnT* also promotes *P. aeruginosa* survival during exposure to ß-lactam and fluoroquinolone antibiotics when grown alongside *E. faecalis* under iron restriction. Likewise, the *P. aeruginosa* c-di-GMP synthase SiaD and the major facilitator transporter MfsC are required for *E. faecalis*-induced antibiotic survival in *P. aeruginosa*, as inactivation of either gene markedly reduced survival following antibiotic challenge across multiple antibiotic classes. Our findings demonstrate that polymicrobial interactions under infection-relevant iron-restricted conditions not only antagonize *P. aeruginosa* growth but also confer an adaptive advantage to the surviving population by altering antibiotic susceptibility.

## RESULTS

### *E. faecalis* and *P. aeruginosa* interactions increase antibiotic tolerance under iron restriction

As previously reported, *E. faecalis* (Ef) strongly antagonizes *P. aeruginosa* (Pa) growth in iron-restricted macrocolony biofilms in a 2,2′-dipyridyl (22D; iron chelator) dose-dependent manner (**Fig. 1A, B**) (33). To determine how this interaction affects antibiotic susceptibility, we challenged 24-hour (h) iron-starved macrocolonies with high doses of an aminopenicillin (ampicillin), a fourth-generation cephalosporin (cefepime), a fluoroquinolone (ciprofloxacin), or a polymyxin (colistin) antibiotic (**Fig. 1A, C**). Percent survival was calculated as (CFU after 24-h antibiotic exposure / CFU after 24-h PBS) × 100, using the matched PBS control of the same biofilm type (single– or mixed-species) performed in parallel; 100% therefore indicates no change relative to PBS, values <100% indicate antibiotic-mediated killing, and values >100% indicate more CFU recovered after antibiotic than after PBS (**Fig. 1C**). The challenge doses greatly exceeded planktonic MICs measured in tryptic soy broth (TSB): ampicillin, 50 mg/mL (∼38-75× MIC); cefepime, 1 mg/mL (∼188-375× MIC); ciprofloxacin, 1 mg/mL (∼1,500-6,000× MIC); colistin, 10 µg/mL (∼7.5-15× MIC), using the MIC ranges obtained for Pa strains PAO1 and PADP6 backgrounds (**Table S4**).

**Figure 1.**
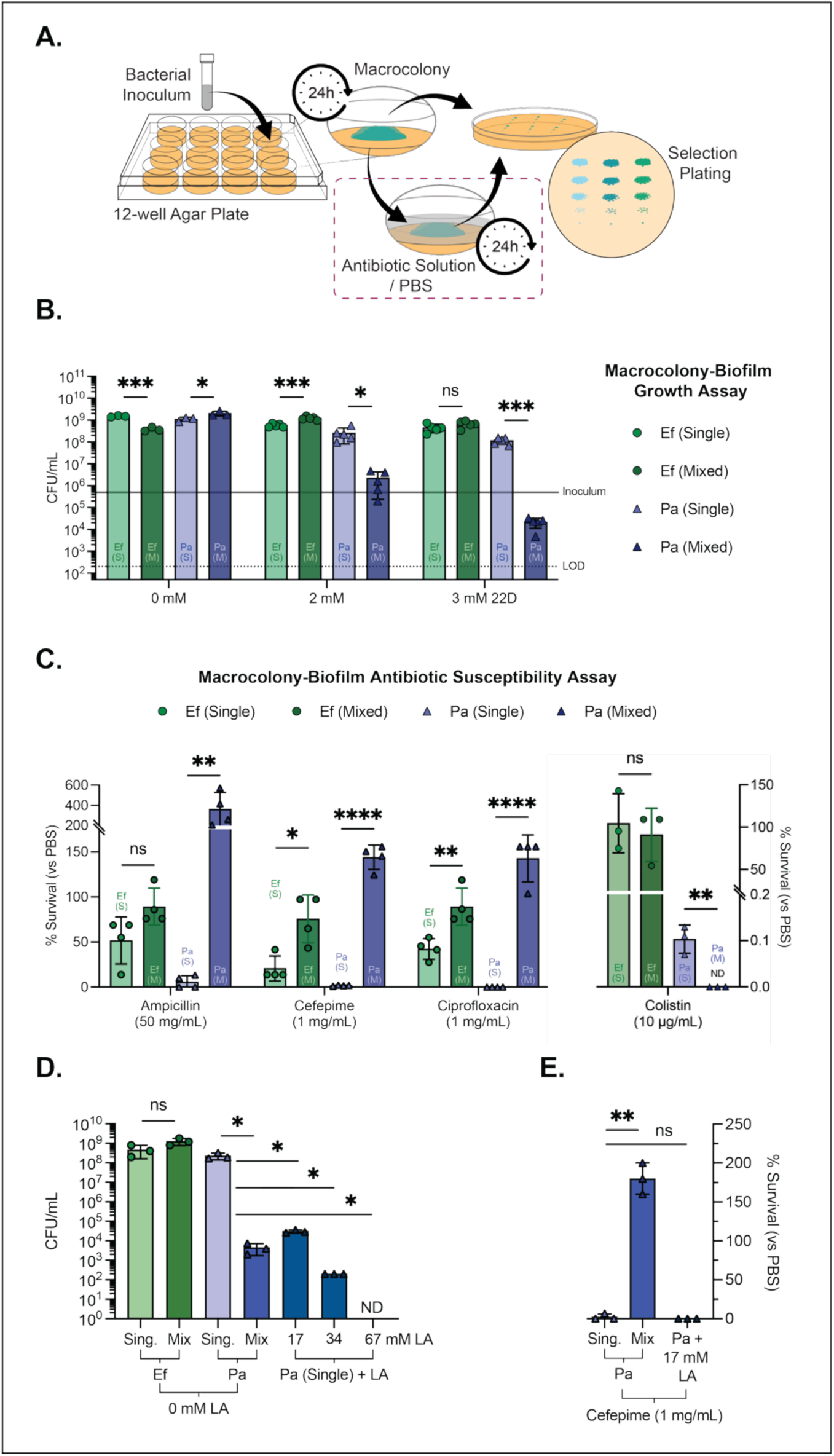
Impact of *E. faecalis* and *P. aeruginosa* interactions on growth and antibiotic tolerance under iron restriction. **(A)** Schematic of the macrocolony growth and antibiotic-challenge assay performed in a 12-well plate format. **(B)** Quantification of bacterial growth as colony-forming units per milliliter (CFU/mL) for *E. faecalis* (Ef; circles, green) and *P. aeruginosa* (Pa; triangles, purple) in single-species (S) and mixed-species (M) macrocolonies grown on iron-restricted (IR) tryptic soy agar (TSB + 10 mM glucose; TSBG) supplemented with the iron chelator 2,2′-dipyridyl (22D; 0, 2, or 3 mM). The solid line denotes the inoculum (5×10^5^ CFU/mL), and the dotted line marks the limit of detection (LOD, 200 CFU/mL). **(C)** Percent survival (%) after 24-h antibiotic challenge (ampicillin 50 mg/mL, cefepime 1 mg/mL, ciprofloxacin 1 mg/mL, or colistin 10 µg/mL) under IR conditions (3 mM 22D TSBG). Each concentration represents tens-to thousands of times the planktonic MIC measured in TSB (see **Table S4**: 38-75× for ampicillin, ∼188-375× for cefepime, ∼1,500-6,000× for ciprofloxacin, and ∼7.5-15× for colistin). Percent survival was calculated as (CFU antibiotic / CFU PBS) × 100, using the matched PBS-treated control of the same biofilm type (single or mixed), where 100% denotes no change relative to PBS, values <100% indicate antibiotic-mediated killing, and >100% indicate increased CFU recovery after antibiotic compared with PBS. Left y-axis % Survival related to ampicillin, cefepime, and ciprofloxacin conditions, while right y-axis % Survival belongs to the colistin treated condition. (**D**) Macrocolony-biofilm growth assay (CFU/mL) on 3 mM 22D (IR) TSBG agar. Ef and Pa in single-IR and mixed-IR macrocolonies are compared with single-IR Pa supplemented with lactic acid (LA; 17, 34, or 67 mM). LA reproduced the growth antagonism observed in mixed-IR (with Ef) macrocolonies (no colonies, ND, detected at 67 mM), but did not confer the increased cefepime survival observed in mixed-IR growth with cefepime challenge (**E**), indicating that acid stress alone is insufficient to recapitulate the Pa antibiotic protective phenotype. Mean ± standard deviation (SD) are shown, and dots indicate biological replicates (*n*). Unpaired t-tests compare single-vs mixed-species biofilms for each antibiotic within a species. Significance is visualized as: ns, not significant; *, P < 0.05; **, P < 0.01; ***, P < 0.001; ****, P < 0.0001. ND not detected (≤ LOD).

In iron-restricted (IR) mixed biofilms with Pa, Ef survival increased compared to single-species biofilms when challenged with cefepime (76% mixed-IR survival vs 21% single-IR) or ciprofloxacin (89% mixed-IR vs 42% single-IR). Ampicillin produced a non-significant reduction in Ef survival in the mixed biofilm (89% mixed-IR vs 52% single-IR), and colistin had little effect in either context (91% mixed-IR vs 105% single-IR). By contrast, IR-Pa survival dramatically increased in the presence of Ef for ampicillin, cefepime, and ciprofloxacin but not after colistin exposure (mixed-IR vs single-IR). Upon ampicillin challenge, Pa monocultures (IR) were reduced to ∼6% survival compared to their PBS control, near the limit of detection (LOD), whereas mixed IR biofilms yielded ∼364% survival, reflecting an ∼3.6-fold increase in CFU compared to the PBS control. Importantly, IR-Ef CFU were not markedly reduced by ampicillin in the mixed IR biofilms (89% mixed-IR survival), indicating that enhanced Pa survival under this condition is not generally explained by antibiotic-mediated elimination of Ef. After cefepime exposure, Pa monocultures (single-IR) were reduced to ∼2% survival, whereas mixed-IR biofilms yielded ∼144% survival relative to the PBS control. Similarly, after challenge with ciprofloxacin, IR monocultures were reduced to undetectable levels, while mixed biofilms yielded ∼143% survival relative to the PBS control, reflecting an ∼1.5-fold increase. This enhanced survival phenotype was not observed with colistin treatment (mixed-IR vs single-IR), which nearly eradicated Pa in both single-and mixed-species IR conditions, while a statistically significant increase in colistin susceptibility was observed in the IR mixed-species condition. Absolute CFU enumerations mirror these patterns (**Fig. S1**). Because Ef-dependent antagonism under iron restriction involves lactic acid production, we asked whether acid stress alone could drive the antibiotic susceptibility phenotype (33). Supplementing Pa monocultures with lactic acid reproduced the growth-inhibitory effect seen in mixed biofilms as previously reported but failed to confer increased survival to cefepime (**Fig. 1D**) (33). Thus, while lactic acid contributes to growth antagonism, it is insufficient to account for the observed increase in Pa survival under these conditions.

Together, these data show that IR growth with Ef generally antagonizes Pa growth (**Fig. 1B**) yet confers a protective advantage to the surviving Pa population when challenged with β-lactams or ciprofloxacin at doses far exceeding planktonic MICs. Colistin is a notable exception, consistent with outer-membrane-targeted killing and the outer-membrane stress signature we describe below. Finally, this unique state cannot be recreated by lactic acid alone (**Fig. 1D-E**), suggesting that factors other than acid stress, likely additional Ef-derived metabolites or signals, are required to elicit antibiotic susceptibility changes in Pa under mixed-species, iron-limited biofilm growth conditions.

### Combined iron limitation and interspecies competition induce a unique transcriptional response in *P. aeruginosa*

RNA sequencing of *P. aeruginosa* (Pa) and *E. faecalis* (Ef) in single– and mixed-species macrocolony biofilms under iron-restricted (IR) and iron-replete (IRp) conditions revealed extensive, context-dependent transcriptional remodeling. Ef showed a minimal transcriptional response to mixed-IR growth with Pa, with only 23 genes upregulated and 35 downregulated (mixed-IR vs single-IR). These differentially expressed genes included modest upregulation of myo-inositol and carbohydrate utilization genes together with selected membrane and transporter loci, while high-affinity phosphate transport and thiamine biosynthesis genes were downregulated (**Table S5**). Notably, many of these loci were altered in the same direction, consistently upregulated or downregulated. When comparing mixed-vs single-species Ef biofilms irrespective of iron restriction, larger amplitudes of expression changes were observed in mixed-IRp vs single-IRp biofilms, indicating that iron restriction reduces the magnitude of the Ef mixed-species response. Evaluating iron restriction alone in Ef (single-IR vs single-IRp), we observed a comparatively greater transcriptional response (471 upregulated, 469 downregulated), enriched for metal and iron acquisition and transport genes such as *adcA, feoB*, and multiple ABC transporters, as well as lactate metabolism genes *larABCD* and inositol catabolism genes (**Table S5**). Under IR conditions, mixed-species biofilm growth with Ef triggered the most extensive changes in Pa, whereby approximately 73% of Pa genes were differentially expressed (mixed-IR vs single-IR) (**Table S5**). Iron restriction alone (single-IR vs single-IRp) induced differential expression for 15% of genes in Pa, and fewer than 7% of Pa genes were differentially expressed in mixed-species biofilms compared to single species growth when iron was not restricted (mixed-IRp vs single-IRp). Principal component analysis (PCA) of Pa single-IR (vs single-IRp), mixed-IR (vs single-IR), and mixed-IRp (vs single-IRp) revealed the distinctiveness of the mixed-IR samples, which separated along PC1 (92.2% variance), indicating the influence of combined iron scarcity and Ef competition on Pa transcription (**Fig. 2A**).

**Figure 2.**
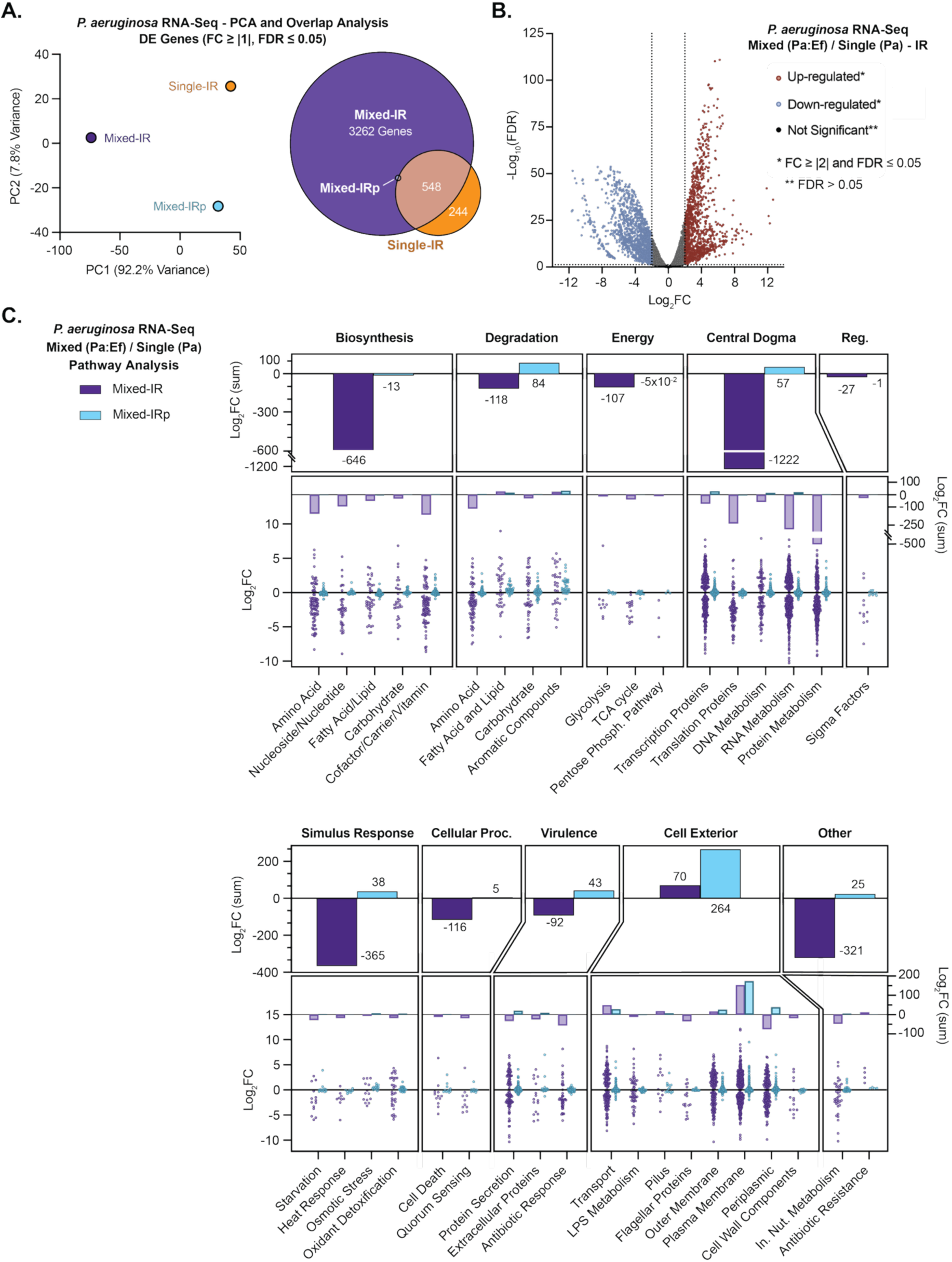
Global transcriptional reprogramming of *P. aeruginosa* under iron-restricted co-culture with *E. faecalis*. (**A**) Principal Component Analysis (PCA) plot of significantly differentially expressed *P. aeruginosa* (Pa) genes with a log2FC ≥ |1| and FDR ≤ 0.05 from RNA sequencing conducted on macrocolony biofilm growth under iron restriction (Single, Iron Restricted), and in mixed-species biofilms with *E. faecalis* (Ef), under both iron restricted (IR) and iron replete (IRp) conditions: single-IR vs single-IRp, mixed-IR vs single-IR, and mixed-IRp vs single-IRp. A Venn diagram depicting these genes is also shown with the number of unique and overlapping genes in each dataset. The mixed-IRp condition contained 2 common genes with mixed-IR and 2 common genes with single-IR. See Table S6 for gene lists. (**B**) Volcano plot visualizing differentially expressed genes in Pa mixed-IR (with Ef) vs single-IR. (**C**) Pathway Analysis (BioCyc) of statistically significant (FDR ≤ 0.05) differentially expressed genes in Pa mixed-IR (with Ef) vs single-IR and mixed-IRp (with Ef) vs single-IRp conditions. Graphs show general pathways as the sum of the log2FC values for the genes in that pathway (top bar graph), followed by the log2FC sum values for the genes in the subpathways (middle bar graph), and finally the individual log2FC values for the genes in each corresponding subpathway (bottom dot plot). See Table S7 for pathway gene lists.

In mixed-IR vs single-IR conditions, Pa exhibited a global adaptive response characterized by broad metabolic downregulation and selective upregulation of nutrient uptake, surface remodeling, and community-associated pathways (**Fig. 2C**). The largest affected groups included cytoplasmic membrane-associated genes, reflecting shifts in membrane composition and transport functionality, with a hallmark of the transcriptomic response in the Pa mixed-IR biofilms being extensive suppression of energetically demanding processes (**Fig. 2C**). Biosynthesis pathways for amino acids, vitamins, fatty acids, and isoprenoids were notably downregulated, as were nucleotide biosynthesis pathways (mean log2FC: purine, –2.53; pyrimidine, –3.47). Central dogma machinery, particularly ribosomal proteins (*rpsU*, –7.36; *rpsR*, –6.90; *rpmH*, –6.85), tRNA processing factors, and sigma factors (*rpoS*, – 4.2; *sigX*, –4.4; *algU*, –1.2), were downregulated, resulting in reduced protein synthesis capacity and growth. Additionally, energy generating pathways, including glycolysis, Entner-Doudoroff, oxidative pentose phosphate pathways, and the TCA cycle, were downregulated. Iron-dependent respiratory complexes similarly decreased, whereas fermentative enzymes and non-iron oxidoreductases were modestly induced, suggesting alternative metabolic routes.

For comparison, single-IR Pa biofilms (vs single-IRp) predominantly activated canonical iron acquisition systems, such as siderophore pyochelin and pyoverdine synthesis genes (e.g. *pchA* to *pchG* gene clusters and *pvdD*, *pvdE*, *pvdH*, *pvdI*, *pvdJ*), as well as endogenous siderophores, heme and ferrous iron OM and/or IM transporter genes (e.g. *fptA*, *fptX*; *fpvA*; *hasR*, *hasAp*, *phuR*; *feoB*) **(Table S5**). Surprisingly, this canonical iron starvation response in Pa was largely suppressed in the presence of Ef (mixed-IR vs single-IR), indicating that Ef significantly interfered or competed with Pa iron-regulated processes. Several genes coding for OM and IM xenosiderophore transporters showed compensatory induction (e.g. *optJ*, *+6.65*; *foxB*, *+5.84*; *fepG*, *+3.18*; *fvbA*, *+2.79*; *fepB*, *+2.44*; *foxA*, *+2.41*; *pfuA*, *+2.30*; *fiuB*, *+2.25*; *PA1909*, *+2.14*; *fecA*, *+1.72*; *pirA*, *+1.64*; *piuA*, *+1.56*).

Despite widespread metabolic downregulation in the mixed-IR condition, Pa pathways related to nutrient uptake and community sensing were selectively activated compared to single-IR growth. Genes for inorganic ion and carbohydrate transporters, particularly ABC and P-type ATPase transporters (*PA1429*, +7.81; *PA3383*, +6.61; *nasA*, +5.40), were strongly induced (mixed-IR vs single-IR), reflecting potential nutrient scavenging adaptations. Efflux systems for heavy metals and toxins (*czcA*, +5.02; *czcB*, +3.78) were also upregulated, possibly to enhance environmental resilience. Additionally, the upregulation of Pa biofilm formation-associated genes was a response uniquely observed under mixed-IR growth (compared to single-IR), reflected in the marked induction of matrix production and adhesion genes (*pelA*, +4.96; *algX*, +4.95; *arr*, +4.83; *cupB1*, +4.23). Upregulation of Pa biosynthesis and cyclic di-GMP signaling genes (*siaD*, +7.03; *pelD*, +4.01; *alg44*, +4.16; *bfiS*, +2.65) in the mixed-IR vs single-IR condition reflects a potential shift toward more sessile, community-oriented lifestyle adaptations. Pa biofilm regulation in the single-IR condition was more variable and limited, further emphasizing the specificity of coordinated biofilm induction in the presence of Ef.

Under mixed-IR growth (vs single-IR), Pa genes associated with lipid A modification via the *arn* operon (*arnBCADT, +1.42 to +8.17*) and the *eptA* gene (+6.23), both known to mediate resistance to antimicrobial peptides, were notably upregulated, consistent with the same proposed outer membrane stress response underpinning the increased colistin sensitivity described in **Figure 1C**. The Pa global regulator *ampR* (+2.51), associated with resistance and stress responses, was similarly upregulated (mixed-IR vs single-IR) suggesting targeted adaptive resistance mechanisms in Pa in response to Ef competition. Traditional antibiotic-response genes, however, were mostly downregulated under the same conditions, emphasizing the specificity of these adaptations. Virulence-associated pathways also showed differential modulation, with general repression of flagella, Type II and III secretion systems, yet selective induction of adhesion factors (**Table S5**). Quorum sensing pathways experienced marked repression of signal synthesis genes (*pqsH*, –4.45), with selective upregulation of specific regulatory genes (*vqsM*, +4.38), indicating a shift toward an energy-conserving, responsive behaviors.

Collectively, mixed iron-restricted growth with Ef, not iron restriction alone, drives a unique and specialized Pa transcriptional response that conserves resources, redirects nutrient acquisition, reinforces the cell envelope, and promotes biofilm formation, in this polymicrobial, iron limited environment.

### Mixed-species-dependent decrease in P. aeruginosa antibiotic susceptibility under iron restriction requires ArnT, PmrA, SiaD, and MfsC and is not reproduced by lactic acid-driven growth inhibition

To define the mechanisms underlying the decreased antibiotic susceptibility of *P. aeruginosa* (Pa) observed in *E. faecalis* (Ef) mixed-species biofilms grown under iron-restricted (IR) conditions (**Fig. 1C**), we assessed antimicrobial susceptibility in Pa isogenic mutants lacking genes identified by RNA sequencing as potential contributors to this phenotype in the mixed-IR vs single-IR dataset (**Table S5**). All susceptibility tests were performed under IR conditions using the same antibiotic concentrations as in **Figure 1C**: ampicillin 50 mg/mL (∼38-75× MIC in TSB), cefepime 1 mg/mL (∼188-375×), ciprofloxacin 1 mg/mL (∼1,500-6,000×), and colistin 10 µg/mL (∼7.5-15×). Percent survival for each condition was calculated relative to the matched PBS control of the same macrocolony biofilm type. Heatmaps of replicate macrocolony assays (z-scores) show that growth with Ef broadly increased Pa survival after β-lactams or ciprofloxacin challenge, whereas colistin remained uniformly bactericidal (**Fig. 3A**). Consistent with Figure 1C, mixed-IR Pa survival increased after cefepime, ampicillin, and ciprofloxacin challenge compared to single-IR, but with no protection against colistin (**Fig. 1C**; **Fig. 3B-C**). Transcriptomic analyses identified induction of Lipid A remodeling, efflux transporters, and biofilm-associated genes in Pa, uniquely in response to combined dual Ef competition and iron limitation. Accordingly, we investigated their contribution to the altered antimicrobial susceptibility phenotype using Pa isogenic mutants lacking ArnT, responsible for the addition of L-Ara4N to Lipid A (40), the regulator of the *arn* operon PmrA (44), the positive regulator of biofilm formation SiaD (45, 46), and the efflux pump component MfsC (47). Loss of *arnT*, *pmrA*, *siaD*, or *mfsC* abolished the increased survival observed for ampicillin, cefepime, and ciprofloxacin, while further amplifying the increased colistin sensitivity observed in mixed-IR vs single-IR growth (**Fig. 3B-E**). Collectively, these data demonstrate that the Ef-dependent decrease in Pa antibiotic susceptibility under iron restriction is an active, genetically encoded and multifactorial response, with at minimum ArnT-mediated lipid A modification together with SiaD-dependent c-di-GMP signaling and the transporter MfsC required, underscoring that mixed species Ef and Pa colocalization under iron limitation engages specific regulatory circuits that subsequently enhance antibiotic survival.

**Figure 3.**
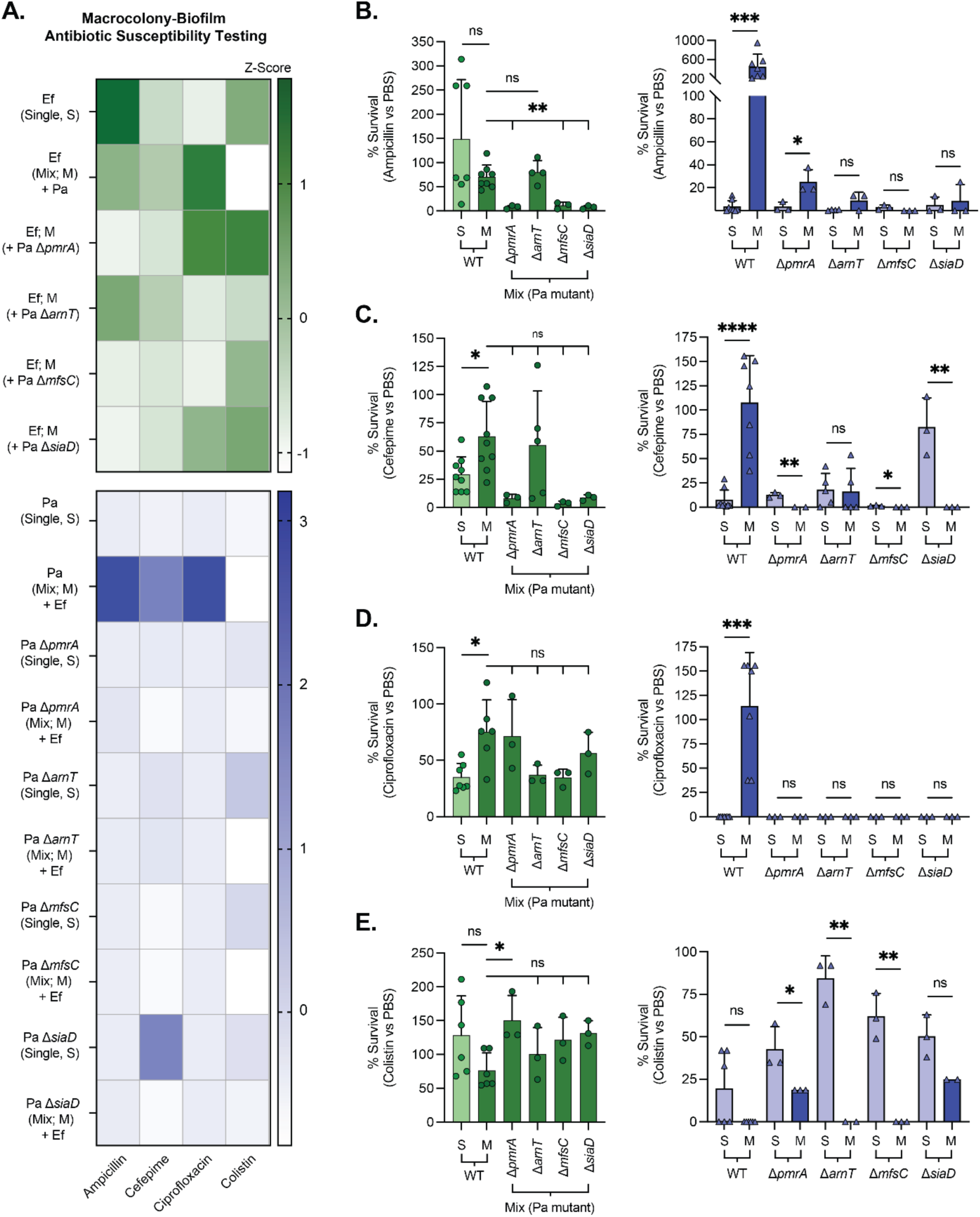
Mixed-species-dependent decrease in *P. aeruginosa* antibiotic susceptibility under iron restriction requires ArnT, PmrA, SiaD, and MfsC. (**A**) Macrocolony-biofilm antibiotic susceptibility heatmaps (iron-restricted medium). Top, *E. faecalis* (Ef; green), shows average percent survival relative to PBS controls for Ef grown alone or mixed with *P. aeruginosa* (Pa) wildtype (WT) or the indicated Pa mutants (Δ*pmrA*, Δ*arnT*, Δ*mfsC*, Δ*siaD*). Bottom, Pa (purple), shows average percent survival for Pa grown alone or with Ef for WT and the indicated Pa mutants. Antibiotic challenges of 24-h macrocolony biofilms for 24-h with ampicillin (50 mg/mL), cefepime (1 mg/mL), ciprofloxacin (1 mg/mL), or colistin (10 µg/mL). Color scale represents z-scores of the mean survival values across macrocolony assays. Mixed-species biofilm growth broadly shifts Pa toward higher survival after β-lactam and ciprofloxacin exposure, whereas colistin sensitivity increases. (**B-E**) Bar graphs show percent survival (antibiotic/PBS) for Ef (left panels, green circles) and Pa (right panels, blue triangles) under the same conditions for (**B**) ampicillin, (**C**) cefepime, (**D**) ciprofloxacin, and (**E**) colistin. Relative to monospecies growth, WT Pa in mixed macrocolonies with Ef exhibited a marked increase in survival to ampicillin, cefepime, and ciprofloxacin, with mixed-species protection lost in Pa mutants for Δ*pmrA* and Δ*arnT* (L-Ara4N lipid-A pathway) and in Δ*siaD* (c-di-GMP synthase) and Δ*mfsC* (major facilitator transporter). Colistin showed minimal or no mixed-species protection for Pa across all genotypes. Ef survival showed greater variation depending on antibiotic and Pa genotype. Bars show means ± SD with individual biological replicates overlaid (n values shown by points; WT single and mixed replicates combined with additional replicates shown in Figure 1C) and survival values were normalized to paired PBS-treated macrocolonies, with statistical significance determined via unpaired two-tailed *t* tests or one-way ANOVA for Ef + Pa mutants; ns, not significant; *, *P*<0.05; **, *P*<0.01; ***, *P*<0.001; ****, *P*<0.0001.

**Figure 4.**
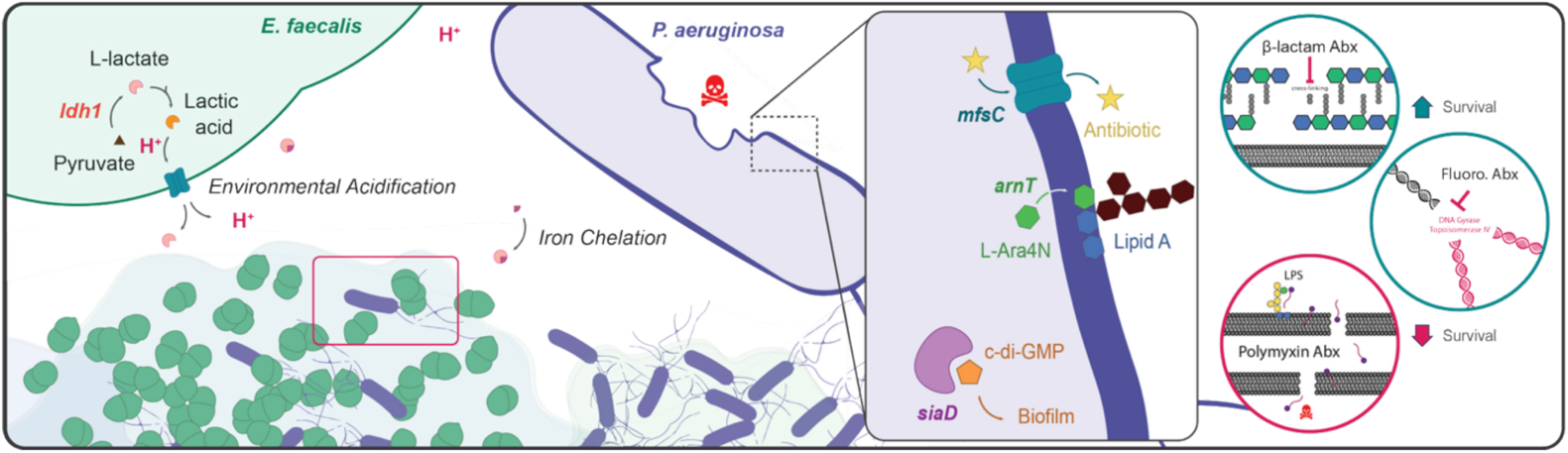
Model of *E. faecalis*-mediated antagonism and antibiotic susceptibility in *P. aeruginosa* under iron restriction. (**A**) Schematic depicting the antagonistic interaction between *E. faecalis* (Ef; green) and *P. aeruginosa* (Pa; purple) in mixed-species macrocolony biofilms grown under iron-restricted (IR) conditions. *E. faecalis* ferments pyruvate to lactic acid via *ldh1*, releasing protons (H⁺) and lactate that acidify the local environment and chelate iron, restricting its availability to *P. aeruginosa* and inhibiting its growth. Within this Ef-altered, low-iron environment, surviving Pa cells exhibit a transcriptionally reprogrammed state characterized by upregulation of *arnT*, *pmrA* (regulator of *arnT*, not shown), *siaD*, and *mfsC*. ArnT and PmrA drive L-Ara4N modification of lipid A, contributing to outer-membrane remodeling and decreased β-lactam susceptibility. SiaD increases intracellular c-di-GMP to promote biofilm-associated tolerance, while MfsC enhances efflux. Together these pathways reduce susceptibility to β-lactams and fluoroquinolones but not to polymyxins (colistin), consistent with envelope-targeted killing. This cooperative antagonism yields a small surviving Pa subpopulation with enhanced antibiotic survival under infection-relevant iron-limiting conditions.

## DISCUSSION

Host-imposed iron restriction is a defining feature of mammalian infection biology, but its impact on bacterial physiology in polymicrobial communities remains largely unexplored (13). Mixed microbial communities, common in wounds, lungs, and urinary tracts, face the dual constraints of nutritional immunity and interspecies competition, and these combined stresses can fundamentally alter microbial dynamics, altering growth, virulence, and antibiotic susceptibility (29, 33, 48, 49). In this study, we define how *Pseudomonas aeruginosa* (Pa) adapts to biofilm growth with *Enterococcus faecalis* (Ef) under iron restriction (IR) and how interactions between these two pathogens, specifically under iron restriction, influence antibiotic susceptibility. Consistent with prior work, Ef strongly antagonized Pa growth in mixed IR macrocolony biofilms, reducing viable Pa by >99% relative to Pa IR monoculture (48). Yet antibiotic challenge of these same mixed species biofilms revealed a Pa subpopulation that displayed significantly altered antibiotic susceptibility at doses far exceeding planktonic MICs. Cooperative effects between Ef and Pa have been noted *in vivo*, where Ef coinfection increased Pa tolerance to β-lactam therapy in a murine pyelonephritis model (29). As with Pa, Ef can facilitate both synergistic and antagonistic interactions with cohabitating bacterial species such as *E. coli* and *S. aureus* depending on environmental context (15, 39, 50–57). Our data indicate that, under combined iron limitation and interspecies competition with Ef, Pa engages a specific adaptive state that complements Ef-mediated growth suppression with enhanced survival following β-lactam and fluoroquinolone antibiotic challenge.

Having profiled both the Ef and Pa transcriptomes under single– and mixed-species conditions with and without iron restriction, we observed interesting and unique transcriptional responses related to each comparative analysis. Mixed iron-replete (IRp) compared to single-IRp growth resulted in minimal transcriptional changes in Ef, with no loci surpassing an absolute log₂FC of 1.5. The small set of altered genes reflect a shift toward increased myo-inositol and carbohydrate utilization and modest activation of stress-associated transporters, coupled with repression of phosphate transport and thiamine biosynthesis pathways. Many of these genes showed larger amplitude changes in mixed-IRp growth compared to mixed-IR biofilms, suggesting that iron restriction significantly attenuates the mixed-species-specific transcriptional response in Ef. Although Ef transcriptomics indicated only modest mixed-IR remodeling, suggesting limited physiological reprogramming prior to antibiotic exposure, these data were collected in the absence of antibiotics. We therefore cannot exclude that antibiotic treatment induces additional post-transcriptional or metabolic changes in Ef not reflected in recovered CFU, which could secondarily influence its antagonistic or protective effects on Pa.

Transcriptomic analysis revealed that Pa in the Ef-mixed-IR condition adopted an expression profile that diverged sharply from both its canonical iron-starvation response in monoculture and from mixed biofilms grown under iron-replete conditions. Rather than the typical induction of iron acquisition systems that dominate the transcriptional response under monoculture iron-limited growth, mixed-IR Pa showed broad transcriptional reprogramming marked by global repression of growth-associated and biosynthetic pathways, including central carbon metabolism, amino acid and nucleotide biosynthesis, and ribosomal protein synthesis, consistent with an energy-conserving, slow-growth state (35, 58). This pattern reflects an active survival strategy rather than passive starvation, where high-cost, iron-intensive processes such as protein synthesis and the TCA cycle were downregulated, while less iron-dependent metabolic routes, including fermentative enzymes and flavodoxin-based oxidoreductases, were selectively induced (59, 60). Such metabolic downshifting has also been shown to enhance antibiotic tolerance by reducing drug target activity and limiting reactive oxygen production and could be helping the surviving Pa cells endure nutrient deprivation while at the same time inducing antibiotic recalcitrance (35, 58, 61–63). In parallel, Pa upregulated transport and stress-response systems under mixed-IR growth, including numerous ion and small molecule transporters, heavy metal efflux components (*czcABC*), and multidrug efflux regulators, mechanisms broadly associated with biofilm-associated antibiotic tolerance (64). Genes involved in cell envelope remodeling were also strongly induced, indicating active reinforcement of the outer membrane in this polymicrobial environment (60, 65). Together, the transcriptional profiling of Pa under dual iron restriction and competition with Ef revealed a resource-conserving, stress-tolerant physiological state that favors long-term survival overgrowth.

One of the strongest Pa transcriptional responses in mixed-IR macrocolonies was the induction of the *arn* operon, which mediates the addition of 4-amino-4-deoxy-L-arabinose (L-Ara4N) to lipid A (43, 65). L-Ara4N decreases the net negative charge of the outer membrane and is a well-established determinant of resistance to cationic antimicrobial peptides (CAMP), such as polymyxins (43). In Pa, *arn* expression is controlled by the PhoPQ and PmrAB two-component systems under divalent-cation limitation (60). Functionally, *arn*-dependent remodeling was necessary for the altered antibiotic susceptibility in Pa observed in mixed-IR biofilms, whereby deletion of *arnT* (L-Ara4N transferase) or *pmrA* eliminated the increased survival of Pa to β-lactams and ciprofloxacin in mixed-IR macrocolonies compared with single-IR growth. This would argue that envelope remodeling contributes broadly to decreased antibiotic susceptibility by reducing drug binding or influx and altering outer-membrane permeability, even for antibiotics that do not directly target LPS (43). While addition of L-Ara4N to lipid A specifically reduces bacteria sensitivity to CAMP by neutralizing the negative charge of its phosphate group, it was recently reported that the arn operon is required for high-level carbapenem tolerance in *Enterobacter cloacae,* likely through enhanced structural integrity of its OM (66). Importantly, *arn* induction was not a default consequence of iron starvation in monoculture, underscoring the polymicrobial triggering of this phenotype (33).

Mixed-IR growth also elevated expression of Pa biofilm-matrix genes (*pel, alg, cup*) and the diguanylate cyclase *siaD*, consistent with increased intracellular c-di-GMP. Elevated c-di-GMP has been shown to promote biofilm formation and increase antibiotic tolerance in Pa by decreasing growth rate, enhancing matrix production, and coordinating stress defenses (67–74). In this model, *siaD* deletion abolished the survival advantage conferred by Ef, indicating that c-di-GMP signaling is required for protection, and the importance of SiaD is congruent with its role in aggregation and surface-associated responses (71). In addition, the major facilitator transporter *mfsC* was necessary for the altered antibiotic phenotypes observed in Pa mixed-IR macrocolonies. Loss of MfsC (Δ*mfsC*) abrogated the increased mixed-species Pa survival to ampicillin, cefepime, or ciprofloxacin. Efflux upregulation is a hallmark of Pa biofilms and antibiotic resistance, contributing by lowering intracellular drug concentration (64). Together, *arn*-dependent LPS remodeling, c-di-GMP-driven biofilm adaptation, and *mfsC*-mediated efflux form a multifactorial defense elicited by Ef under iron restriction. In addition to matrix-associated effects, polysaccharides themselves can provide immediate protection by limiting diffusion and binding antibiotics (68, 75). Our mixed-IR transcriptome showed coordinated induction of matrix loci, consistent with a role for the extracellular matrix in fast-acting defense (75). While we did not directly quantify matrix composition here, the genetic requirement for *siaD* and the induction of *pel*, *alg*, and *cup* genes argue that matrix-mediated barriers act in concert with *arn*-dependent envelope remodeling and efflux to protect a surviving antagonized Pa subpopulation (64, 68).

In Pa monoculture, iron starvation typically induces siderophore production and iron-uptake systems, such as pyoverdine/pyochelin biosynthesis, receptors, and heme uptake (38, 76–79). Counterintuitively, many of these systems were downregulated in Pa during mixed-IR growth despite severe iron limitation. Ef is known to undergo fermentative metabolism under iron restriction, exporting L-lactate that acidifies the milieu and chelates iron, and we previously showed an Ef Δ*ldh1* mutant, defective in lactic-acid production, loses the ability to antagonize Pa growth (33). These effects potentially render siderophore-mediated iron scavenging ineffective for Pa, favoring energy conservation and stress tolerance over more futile iron-acquisition efforts (33). Thus, Ef reshapes the microenvironment to impose a form of nutritional interference that suppresses the canonical Pa iron-uptake response. Whether Pa additionally detects contact-dependent or metabolite signals from Ef that directly repress iron-acquisition regulons remains to be determined. Importantly, we showed in this study that lactic acid alone was unable to reproduce the increased antibiotic survival phenotype in Pa. Exogenous lactic acid recapitulated Pa growth inhibition under IR but failed to confer increased survival to cefepime, indicating that additional cues generated only during cohabitation with Ef are required for the protective state (33). These cues could potentially include contact-dependent signaling, peptide or small molecule effectors, or redox or ionic changes that together drive the protective state (17, 80–87). Distinguishing which signals are necessary and sufficient will be important in future work to clarify how Pa prioritizes survival pathways over iron scavenging during polymicrobial stress.

The mixed-IR transcriptome of Pa also featured strong induction of *ampR*, a LysR-type global regulator controlling expression of over 500 genes, including β-lactamase (*ampC*), efflux pumps, quorum sensing, and virulence factors (88). Although AmpR can repress biofilm formation under some conditions, our data showed simultaneous upregulation of biofilm gene sets, implying that c-di-GMP signaling and iron-stress networks may override or redirect AmpR outputs in this dual-stress setting (88). Because AmpR integrates cell wall and envelope stress signals, its induction may help coordinate the *arn*/efflux arm of the response, while other regulators such as c-di-GMP circuits or stringent response drive matrix and growth arrest (61, 88). Furthermore, the reduced growth or cell-cycle arrest of *P. aeruginosa* in mixed-species biofilm under iron restriction is likely to affect AmpR transcription factor activity since specific peptidoglycan degradation products imported in the cytoplasm act as activator of AmpR, while a peptidoglycan biosynthetic precursor promotes repression (89). Testing *ampR* mutants in this model will be useful to clarify whether AmpR directly contributes to the tolerance phenotype or reflects broader envelope stress.

Standard monoculture susceptibility tests would not have predicted the highly altered antibiotic susceptibility states observed for Pa survivors in mixed-species IR biofilms. Polymicrobial biofilms are often more tolerant than monospecies biofilms, reflecting community-level defenses (7, 68), however, in our assays, this broad increase in Pa survival did not extend to colistin, which remained generally effective in both mixed– and single-IR contexts. Although L-Ara4N addition can decrease polymyxin susceptibility (43, 60), the extent and kinetics of *arn*-dependent modification here appear insufficient to protect against colistin membrane-disruptive action or reflects an ability to overwhelm a partial defense. Notably, the genetic determinants tested in this study, *arnT*, *pmrA*, *siaD*, and *mfsC*, were selected because they were uniquely upregulated during mixed-IR growth. Their deletion nonetheless altered colistin susceptibility in single-IR monocultures, suggesting that the observed hypersensitivity in the single-IR mutants reflects secondary effects, potentially such as envelope instability or pleiotropic stress responses, rather than loss of the mixed-IR-specific mechanism itself. This class specificity suggests that certain antibiotics can still circumvent the induced defenses, which may help guide therapy in polymicrobial infections where β-lactams or fluoroquinolones underperform.

Finally, we acknowledge that the macrocolony biofilm model on defined medium used in this study is a simplified approximation of an iron-restricted chronic infection niche, where certain complexities associated with the spatial, immunologic, and nutrient complexity of chronic infections are not represented. Nevertheless, the insights gained here are likely relevant to infections where Pa and Ef co-infect, or in subpopulations where local iron limitation and acids from fermentative metabolism are concentrated. The relevant pathways we identify, *arn*-dependent lipid A modification, SiaD-mediated c-di-GMP signaling, and *mfsC*-linked efflux, offer potential points of therapeutic leverage. Agents that inhibit c-di-GMP signaling or diguanylate cyclases could limit matrix-associated tolerance (68), while compounds that block L-Ara4N addition to lipid A such as by targeting PmrAB/ArnT or disrupting outer-membrane charge, may resensitize Pa (43, 60, 65, 90). Strategies that reduce Ef lactate production or neutralize acidification may also prevent Pa from entering this tolerant state (33). More broadly, our findings argue that managing polymicrobial infections may require interventions that also disrupt beneficial interactions between pathogens, in addition to directly targeting the organisms themselves (12, 68, 91). While the combination of host-imposed iron limitation and interspecies competition drives a distinct survival program in Pa that is not revealed by single-stressor or single-species assays, mixed-species IR biofilms with Ef suppressed Pa growth-related metabolism, downregulating canonical iron acquisition, and engaging a multifactorial defense consisting of *arn*-dependent outer-membrane remodeling, c-di-GMP–driven biofilm adaptation, and *mfsC*-linked efflux. This state produced a cooperative protection against β-lactams and ciprofloxacin while leaving Pa vulnerable to colistin. Appreciating and therapeutically targeting such context-specific adaptations will be crucial for managing polymicrobial infections where conventional monotherapy more frequently fails.

## MATERIALS AND METHODS

### Bacterial strains and growth conditions

Bacterial strains used are described in Table S1. Strains were grown statically at 37 °C from single colonies on tryptic soy broth (TSB; Sigma-Aldrich, USA) agar (Difco BD, USA) in 6 mL of TSB. Overnight cultures were harvested at 4,300 × *g* for 5 minutes, washed with sterile phosphate-buffered saline (PBS; Sigma-Aldrich, USA), resuspended in PBS and then adjusted to 1 × 10^8^ CFU/mL for further use. When required, bacterial selection and differentiation was performed on TSB agar supplemented with either 25 µg/mL rifampicin (Sigma-Aldrich, USA) for *E. faecalis* or 100 µg/mL ampicillin (A0839 BioChemica; AppliChem GmbH, Darmstadt, Germany) for *P. aeruginosa*. For iron-restricted media, TSB agar supplemented with 10 mM glucose (TSBG) (Sigma-Aldrich, USA) was used with a final concentration of 3 mM 2,2’-dipyridyl (22D) (D216305; Sigma-Aldrich, USA), prepared from stock at 100 mg/mL in ethanol.

### *P. aeruginosa* strain construction

Gene deletions were constructed by two-step allelic exchange using vector pEXG2 (92). Briefly, pEXG2 vectors carrying the desired mutation were introduced in P. aeruginosa by conjugation from a the diaminopimelic acid (DAP) E. coli strain WM3064. Merodiploids were selected on gentamicin, sucrose counter-selection was used to select for double crossovers, and mutant carrying the desired mutation were screened by PCR. The primers used to generate deletion alleles of mfsC, pmrA, and siaD with upstream and downstream flanking regions of the target locus in the PADP6 genome by overlap extension PCR are listed in Table S3. These deletion alleles were designed to retain the start, stop, and 5 to 7 codons to avoid potential polar effect. Plasmid inserts were verified by Sanger sequencing. The pΔarnT vector used to generate the arnT deletion was described previously (93).

### MIC determinations

MICs were determined in Tryptic Soy Broth Glucose (TSBG) at 37 °C by broth microdilution in accordance with Clinical and Laboratory Standards Institute (CLSI) guidelines using inoculum from static culture in TSB (94). All MICs were determined at least 4 times independently using the following concentration ranges (µg/mL): ampicillin, 20.8 to 1333.3; cefepime, 0.167 to 10.667; ciprofloxacin, 0.042 to 2.667; colistin, 0.167 to 10.667.

### Macrocolony biofilm assays

Macrocolony biofilm growth assays were performed by adding 1.5 mL of 3 mM 22D TSBG agar to wells of a sterile 12-well plate (BDAA353043; Corning, USA). Bacterial inocula under single species (5 µL inoculum) or mixed species (10 µL, 1:1 mixture) conditions were prepared as described above and spotted with one replicate per well onto the prepared agar. After drying, the plates were incubated at 37 °C for 24 hours before being harvested by washing twice with 1 mL of PBS. Harvested macrocolonies were then serially diluted in a 96-well plate and 5 µL plated on selection media as described above. After 24-hour incubation at 37 °C, CFUs were enumerated to determine the CFU/mL. Results were plotted and analyzed using GraphPad Prism (v10.6.1; GraphPad Software, Boston, MA, USA).

### Macrocolony RNA extraction

RNA-seq was performed on macrocolonies grown as described above after 24-hour incubation on TSBG agar with or without 2 or 3 mM 22D in triplicates. The macrocolonies were harvested by washing twice with 0.7 mL of 1:2 PBS:bacterial RNAprotect (Qiagen, Germany), then incubating at room temperature for 5 minutes. The samples were then centrifuged at 5,000 x *g* for 10 minutes, the supernatant was discarded, and the pellet resuspended in 200 µL of lysis solution (1x TE, 20 mg/mL lysozyme (A3711,0001, Applichem), 100 U/mL Ready-Lyse (E0057-D3, Biosearch Technologies)) before the addition of Proteinase K (15 µL; PA4850, Sigma-Aldrich). The samples were then incubated at 37 °C for 30 minutes on a heat block before the addition of 1 mL of TRIzol reagent (Invitrogen, USA). At this point, samples could either be stored at –80 °C or further processed. RNA extraction was performed by taking the samples in TRIzol reagent and adding 200 µL of chloroform per 1 mL TRIzol. Tubes were vigorously shaken for 15 seconds, then incubated for 3 minutes at room temperature. Samples were then centrifuged at 12,000 × *g* for 15 minutes at 4 °C and the top aqueous phase transferred to a new 1.5 mL tube. Ethanol (80%) was added (550 µL EtOH/700 µL aqueous phase) and transferred, 700 µL at a time to an RNeasy Mini Spin column (71404, Qiagen) for centrifugation at 8,000 × *g* for 15 seconds. Buffer RW1 was added (350 µL) and centrifugation repeated. A prepared DNase mixture (10 µL DNase I with 70 µL Buffer RDD per sample, 79254, Qiagen) was added to the columns (80 µL) for on-column DNase digestion and incubated at room temperature for 15 minutes. Then 350 µL of Buffer RW1 was added, followed by centrifugation, the addition of 500 µL of Buffer RPE with centrifugation, and finally 500 µL of Buffer RPE and centrifugation for 2 minutes at 8,000 × *g*. RNA was finally eluted into 45 µL of RNase-free water, repeating with the initial eluate. A second DNase digestion was performed with RQ1 DNAse (8U/10µg RNA, M6101, Promega) in presence of RNasin Plus RNase Inhibitor (8U/10µg RNA, N2611, Promega) for 30 minutes at 37°C and stopped with RQ1-DNAse stop solution. After addition of RLT buffer supplemented with β-mercaptoethanol, RNA cleanup was performed using an RNeasy Mini Spin column as described above. Purified RNA was quantified using a spectrophotometer (Ultrospec 3100 pro, Amersham Biosciences) and RNA quality was assessed using a 2100 Bioanalyzer system (Agilent).

### RNA-sequencing and analysis

RNA was sequenced by the iGE3 Genomics Platform of the University of Geneva as paired-end 100-bp reads on a NovaSeq 6000 instrument (Illumina) after library preparation using the Ribo-Zero Plus rRNA Depletion kit (Illumina). The amount of the libraries prepared from mixed-species samples under restriction was increased 10 times in the sequenced pool to account for the low *P. aeruginosa* CFU numbers in theses sample. RNA sequencing data as fastq.gz files were processed using a custom Python pipeline, where paired-end reads were aligned to the reference genome (*E. faecalis* OG1RF, NCBI Accession No. CP002621; *P. aeruginosa* PAO1, NCBI Accession No. NC_002516) using BWA-MEM (version 0.7.18), the resulting BAM files sorted using Samtools (version 1.20), and gene-level quantification was performed with HTSeq-count (version 2.0.7) using coding sequences (CDS) and locus tags as identifiers, with a minimum alignment quality threshold of 10. Raw feature counts from all samples were aggregated into a single CSV file for downstream analysis. Raw feature count data were then imported into R (version 4.3.1) with RStudio (version 2023.12.1+402) and preprocessed for differential gene expression analysis using edgeR (version 4.0.16) (95). The data were filtered to remove genes with low expression levels, and normalization was performed to account for differences in library sizes. For each condition, data were modeled using a generalized linear model (GLM) approach. Dispersion was estimated, and differential expression was assessed using a likelihood ratio test to compare expression between groups. Gene-level results were then annotated with functional information, and the top differentially expressed genes were extracted and saved for further analysis. For pathway analysis, clusterProfiler (version 4.12.6) was used to perform gene set enrichment analysis using KEGG pathways and identifiers (gseKEGG).

### Macrocolony antibiotic susceptibility assays

Macrocolony antibiotic susceptibility testing was performed as described for macrocolony growth assays, with the addition of 24-hour antibiotic exposure before harvesting. Macrocolonies grown for 24 hours were subjected to 1 mL of antibiotic solution in PBS, or a PBS control, added gently to the well to prevent disrupting the biofilm. After an additional 24-hour incubation at 37 °C with antibiotic, the solution was homogenized in the well on top of the agar, removed to a microcentrifuge tube, and the well washed with an additional 1 mL of PBS. The harvested sample was centrifuged at max speed for 5 minutes, the supernatant discarded, pellet washed with 1 mL of PBS, centrifugation repeated, and then finally resuspended in 1 mL of PBS for dilution plating as previously described. The antibiotic-treated condition was normalized to the PBS control to determine the percent survival following antibiotic treatment. Antibiotics used with preparation details can be found in Table S2. Results were visualized and analyzed using GraphPad Prism (v10.6.1; GraphPad Software, Boston, MA, USA).

### Statistical analysis and software used

Statistical analyses were performed using GraphPad Prism (v10.6.1; GraphPad Software, Boston, MA, USA) with specific tests described in the relevant figure legends. Figures were prepared using Adobe Illustrator (version 26.3.1). RNA processing was performed using custom python scripts in Visual Studio Code (version 1.87.0) with downstream analysis performed using edgeR (version 4.0.16) with R (version 4.3.1) in RStudio (version 2023.12.1+402).

### Data availability

RNA sequencing data have been deposited in the National Center for Biotechnology Information (NCBI) Sequence Read Archive (SRA) database under BioProject PRJNA1365690.

## Acknowledgements.

We are grateful to colleagues at the UNIGE iGE3 Sequencing Facility for performing library preparation and RNA sequencing. This work was primarily supported by funding from the Fondation privée des HUG to K.A.K and P.H.V. (RC08-20). This was also supported by funding to K.A.K. from the Société Académique de Genève and from the Swiss National Science Foundation (310030_219227), as well as by the Singapore Centre for Environmental and Life Science Engineering (SCELSE), whose research is supported by the National Research Foundation Singapore, Ministry of Education, to Nanyang Technological University and the National University of Singapore under its Research Centre of Excellence program. A.P. was supported by a postdoctoral fellowship from the Regional Government of Galicia, Spain (Reference: ED6481B-2023/117).

## Author Contributions

Caleb M Anderson – experimental design, manuscript, performed experiments, data analysis

Yves Mattenberger – experimental design, strain construction performed experiments, data analysis

Ana Parga – performed experiments, experimental design, data analysis

Heidi Portalier – performed experiments, data analysis

Casandra Ai Zhu Tan – performed experiments and data analysis

Maria Esteban Henao – strain construction, performed experiments

Patrick H Viollier – manuscript, funding acquisition, experimental design, data analysis

Kimberly A Kline – manuscript, funding acquisition, experimental design, data analysis

